# Compensating complete loss of signal recognition particle during co-translational protein targeting by the translation speed and accuracy

**DOI:** 10.1101/2021.01.06.425660

**Authors:** Liuqun Zhao, Gang Fu, Yanyan Cui, Zixiang Xu, Tao Cai, Dawei Zhang

**Affiliations:** Tianjin Institute of Industrial Biotechnology, Chinese Academy of Sciences, Tianjin, China; Key Laboratory of Systems Microbial Biotechnology, Chinese Academy of Sciences, Tianjin, China; National Engineering Laboratory for Industrial Enzymes, Chinese Academy of Sciences, Tianjin, China; University of Chinese Academy of Sciences, Beijing, China

**Keywords:** signal recognition particle, suppressor screening, translational control, inner membrane protein, co-translational protein targeting

## Abstract

Signal recognition particle (SRP) is critical for delivering co-translational proteins to the bacterial inner membrane. Previously, we identified SRP suppressors in *Escherichia coli* that inhibit translation initiation and elongation, which provided insights into the mechanism of bypassing the requirement of SRP. Suppressor mutations tended to be located in regions that govern protein translation under evolutionary pressure. To test this hypothesis, we re-executed the suppressor screening of SRP. Here we isolated a novel SRP suppressor mutation located in the Shine-Dalgarno sequence of the S10 operon, which partially offset the targeting defects of SRP-dependent proteins. We found that the suppressor mutation decreased the protein translation rate, which extended the time window of protein targeting. This increased the possibility of the correct localization of inner membrane proteins. Furthermore, the fidelity of translation was decreased in suppressor cells, suggesting that the quality control of translation was inactivated to provide an advantage in tolerating toxicity caused by the loss of SRP. Our results demonstrated that the inefficient protein targeting due to SRP deletion can be rescued through modulating translational speed and accuracy.

## INTRODUCTION

The signal recognition particle (SRP) is a highly conserved ribonucleoprotein complex that is involved in co-translational targeting of the ribosome-nascent chain complex to the endoplasmic reticulum of eukaryotes or the inner membrane of prokaryotes (Pool, 2005; Walter and Johnson, 1994). Although the size and composition of SRP are variable in different species, the key subunits of SRP are evolutionarily conserved (Saraogi and Shan, 2014). The *Escherichia coli* SRP is much simpler than that in eukaryotes, as it contains an essential and highly conserved subunit called the fifty-four homologue (Ffh), which is homologous to mammalian SRP54, and a small stable 4.5S RNA, which is homologous to domain IV in the mammalian 7S SRP RNA (Bernstein et al., 1993; Powers and Walter, 1997; Regalia et al., 2002). The *E. coli* SRP components can replace their mammalian homologues to mediate efficient co-translational protein targeting of mammalian proteins (Bernstein et al., 1993; Powers and Walter, 1997). This suggests that the subunit SRP54 and domain IV of the 7S SRP RNA form the core elements of SRP, and SRP is remarkably conserved from bacteria to mammals. SRP is primarily responsible for delivering inner membrane proteins (Ulbrandt et al., 1997a; Zhang and Shan, 2014). It recognizes hydrophobic transmembrane domains or signal sequences when they emerge from the ribosome exit tunnel (Ng et al., 1996). Furthermore, the nascent polypeptides must be targeted by SRP in a limited time window before they lose their competence due to aberrant aggregation (Flanagan et al., 2003; Siegel and Walter, 1998). Given the crowded cellular environment, it is challenging to correctly translocate newly synthesized proteins from the cytosol to the membrane.

SRP is generally essential in all three kingdoms of life (Akopian et al., 2013; Egea et al., 2005), except for the eukaryote *Saccharomyces cerevisiae* (Mutka and Walter, 2001) and prokaryotes *Streptococcus mutans* (Kremer et al., 2001) and *E. coli* (Zhao et al., 2021). In these cases, the global repression of protein synthesis is associated with the loss of SRP (Hasona et al., 2007; Mutka and Walter, 2001; Zhao et al., 2021). However, the precise mechanism of this tolerance remains poorly understood (Hasona et al., 2007; Kremer et al., 2001; Mutka and Walter, 2001; Zhao et al., 2021). One of the best-understood mechanisms is slowing the translation elongation rate that extends the time window for targeting translating ribosomes. Previous studies have demonstrated that slowing translation elongation speed contributes to improving protein folding and targeting (Sherman and Qian, 2013; Siller et al., 2010; Zhang and Shan, 2012). In co-translational translocation, protein folding and targeting are inherently coupled to translation elongation (du Plessis et al., 2011). The kinetic competition between protein translation elongation and targeting modulates the efficiency of the co-translational targeting pathway (Chartron et al., 2016; Zhang and Shan, 2012). In eukaryotes, the Alu domain of SRP arrests nascent chain translation elongation during its targeting, which is thought to pause translation elongation until the targeting is completed (Mason et al., 2000; Walter and Blobel, 1981). Furthermore, the Shine-Dalgarno (SD)-like sequence pauses translation before the second transmembrane domain exposed in *E. coli*, which facilitates the proper folding and targeting of membrane proteins (Fluman et al., 2014). Thus, the translation elongation is at the center of protein folding and targeting.

In our previous study, we identified SRP suppressors: two translation initiation factors IF2 and IF3, and a ribosomal protein RS3. The suppressor mutations decreased the translation initiation and elongation rate (Zhao et al., 2021). There are two possible explanations for the slowing translation elongation rate. First, it is possible that suppressor mutations directly inhibit the translation initiation and then decrease the translation elongation rate because the translation initiation rate is closely correlated with the elongation rate (Riba et al., 2019; Zhao et al., 2021). Alternatively, mutations that suppress the lack of SRP can directly affect the translation elongation rate, rather than indirectly decrease the rate of translation initiation. The ribosome is a hub in protein translation, which can directly modulate translation elongation (Pechmann et al., 2013; Sherman and Qian, 2013). Here, we isolated an SRP suppressor located at the SD sequence of ribosome S10 operon that could affect ribosome biogenesis. We addressed how the suppressor regulated the translation process and alleviated the fitness loss. Our results suggested that this mutation suppressed the loss of SRP, although the cell growth was severely inhibited and the targeting of SRP-dependent proteins was not completely compensated. This SRP suppressor reduced translation rate. Moreover, the translation initiation and elongation fidelity were decreased to improve cell viability. Overall, our results showed that mutations in the SD sequence of ribosome S10 operon contributed to protecting cells from lethal damages caused by the loss of SRP.

## MATERIALS AND METHODS

### Bacterial Strains, Plasmids and Media

All bacterial strains, plasmids and primers used in this study are listed in **Supplementary Table S1**. *E. coli* K-12 MG1655 derivative strains were grown either in LB medium or on LB agar at the indicated temperature. *E. coli* HDB51 strain in which the expression of Ffh is under the control of an arabinose inducible promoter, were grown in LB medium containing 0.2% arabinose (Lee and Bernstein, 2001). The SRP suppressor strain MY1901 was isolated and validated as previously described (Zhao et al., 2021). The antibiotics kanamycin, gentamycin, ampicillin, and chloramphenicol were used at a concentration of 50, 10, 100, and 200 μg ml^-1^, respectively. The *lac*, *trc* and *tac* promoters were induced with 0.02 mM isopropyl-β-D-1-thiogalactopyranoside (IPTG). The *araBAD* promoter was induced with 0.2% arabinose.

### Polysome Analysis, Cell Ultrastructure and Proteomic Analysis

Strains were grown at 37°C at the early-exponential growth in LB medium and harvested by centrifugation (Zhao et al., 2021). Polysome analysis was performed as described (Zhao et al., 2021). The gradients were first extracted with a Piston Gradient Fractionator (Quan et al., 2005), and then their UV spectra were monitored by the ÄKTA equipment (Malecki et al., 2014). The scanning electron microscopy (SEM) and transmission electron microscopy (TEM) analyses were carried out as described previously (Zhao et al., 2021). The tested cells were randomly selected. The whole-cell lysates and inner membrane proteins of strains MY1901 and SRP^−^ were isolated and analyzed by liquid chromatography-tandem mass spectrometry (LC-MS/MS) as described (Tsolis and Economou, 2017; Wisniewski et al., 2009; Wisniewski and Mann, 2012; Zhao et al., 2021), with modifications. For seed culture, the strain MY1901 or HDB51was inoculated in LB medium at 37°C overnight and the strain HDB51 was grown under the addition of 0.2% arabinose. The overnight culture of MY1901 was diluted into the fresh LB medium with an initial OD_600_ of 0.02-0.03. Strain MY1901 was harvested in the mid-exponential phase. The overnight culture of HDB51 was washed for three times with fresh LB medium and then incubated in LB with addition of 0.2% glucose with an initial OD_600_ of 0.03-0.04 for several hours, yielding the SRP^−^ strain. This strain was harvested when entering the stationary phase. Three sample replicates were prepared by performing collection of cells from independent cultures. We used the Filter-Aided Sample Preparation (FASP) strategy for proteome analysis (Wisniewski et al., 2009). The inner membrane fraction was separated by sucrose gradient (Tsolis and Economou, 2017). Then, the inner membrane was chemically treated by Na_2_CO_3_ and KCl to remove peripherally associated proteins (Zhao et al., 2021). The MS samples of inner membrane proteins were prepared by surface proteolysis (Tsolis and Economou, 2017). The sample was characterized by sequential window acquisition of all theoretical spectra (SWATH) analysis (Jylhä et al., 2018). The LC-MS/MS analysis was performed according to the previous study (Zhao et al., 2021). The obtained data were normalized by the median scale normalization (MedScale) method (Callister et al., 2006).

### β-galactosidase Assay

The β-galactosidase activity was assayed as previously described (Miller, 1972; O’Connor et al., 1997). Cells were grown in LB medium supplemented with 50 μg ml^-1^ kanamycin at 37°C. Protein induction was performed when MG1655Δ*lacZ* and MY1901Δ*lacZ* were grown to OD_600_ of 0.6-0.8 and 0.4-0.5, respectively. Two hours later the cultures were harvested and assayed. β-galactosidase activity from the plasmid encoding wild-type LacZ was used for normalization.

### Measurements of Translation Efficiency and Initiation Fidelity

To determine the effects of mutated SD sequence on protein translation efficiency, we constructed plasmids carrying the *gfp* gene with the wild-type SD and suppressor mutation SD* under control of the promoter P*_S10_* or P*_araBAD_*. To analyze the translation initiation fidelity, a set of GFP variants were generated in which the start codon (AUG) and the initiator tRNA were replaced with different start codons and non-initiator tRNAs codons, respectively. Several non-AUG start codons (GUG, UUG, AUA, and AUC) and non-initiator tRNAs codons (UAG: CUA, CAC: GUG, and UAC: GUA) were used as potential start codons and initiator tRNAs codons, respectively. *E. coli* cells were grown in 300 μl of LB medium with necessary antibiotics in a 96-well deep well culture plate at 37°C overnight in a stationary phase (MG1655Δ*lacZ*, OD_600_>3.0; MY1901Δ*lacZ*, OD_600_>1.0) followed by transferring and cellular fluorescence measurements (Hecht et al., 2017). For detecting the effects of Shine-Dalgarno (SD) sequence on translation efficiency and the fidelity of translation initiation, the strains containing empty vectors pJH30, pJH31, and pTrc99K lacking a reporter gene were used as controls. OD_600_ was measured to estimate culture density, followed by fluorescence (excitation=488 nm, emission=520 nm). Assays were carried out from at least three independent colonies. For the initiation fidelity assay, the fluorescence intensity of strains containing the plasmid encoding mutated GFP was normalized against the fluorescence intensity of wild-type or suppressor strains containing plasmid-encoded wild-type GFP.

### Measurements of Translation Rates

Translation elongation rates were measured as described previously (Dai et al., 2018; Dai et al., 2016; Zhu et al., 2016). For seed culture, MG1655Δ*lacZ* and MY1901Δ*lacZ*Δ*cat* cells were cultured in LB medium at 37°C for several hours, then cultures were collected and washed with fresh MOPS medium. To improve the growth of the strains MG1655Δ*lacZ* and MY1901Δ*lacZ*Δ*cat*, they were grown in rich Glucose + cAA (0.2% glucose + 0.2% casamino acids) MOPS medium overnight as the pre-culture. The experiment culture was performed with an initial OD_600_ of 0.04-0.05. The subsequent collection and measurement methods were performed as previously described (Zhao et al., 2021). The translation elongation rate was measured based on the LacZα induction assay.

The translation time of the first newly synthesized LacZα, T_α_, was estimated by measuring the LacZα induction kinetics. The translation time of the first newly synthesized LacZα fused protein (FusA-LacZα or MsbA-LacZα), T_total_, was obtained by the Schleif plot of the induction curve. The initiation time, T_init_, equals T_α_–{90/[L/(T_total_–T_α_)]}, where 90 is the 90 aa LacZα fragment and L is the length of the LacZα fusion protein (containing 10 aa linker). The translation elongation rate equals (L+90)/(T_total_–T_init_). When MY1901Δ*lacZ*Δ*cat* grew in Glucose + cAA medium, the growth rate was about 0.6 h^-1^. To eliminate the cell growth effect on translation elongation rate, the wild-type strain MG1655Δ*lacZ* was grown in Glycerol + NH_4_Cl (0.2% glycerol + 10 mM NH_4_Cl) MOPS medium and the growth rate was similar to 0.6 h^-1^. The measurement of the elongation rate of MG1655Δ*lacZ* with the growth rate of 0.6 h^-1^ was performed similarly to that of MY1901 Δ*lacZ*Δ*cat*. Translation initiation rate was estimated by a computational model homogeneous ribosome flow model (HRFM) (Margaliot and Tuller, 2012; Zarai et al., 2013; Zarai et al., 2014). Based on the measured translation rate and elongation rate, the initiation rate can be calculated (Zhao et al., 2021).

### Protein Targeting Assay *in vivo*

The biotinylation of proteins has been successfully applied to SRP-dependent protein targeting in *vivo* (Jander et al., 1996; Zhang and Shan, 2012). *E. coli* enzyme biotin ligase (BirA) can specially ligate biotin to a 15-amino acid peptide (GLNDIFEAQKIEWHE) termed the Avi-tag (Chen et al., 2005). The BirA and Avi-tagged proteins were co-expressed for biotinylation. Cells were co-transformed with recombinant vectors p15A-birA and pJH29-EspP/FtsQ/LacZ-Avi and grown overnight at 37°C. Then overnight cultures were washed and diluted into 30 mL fresh LB medium at an initial OD_600_ of 0.02. For HDB51, the overnight culture grown in LB medium with 0.2% arabinose was washed and diluted into 30 mL fresh LB medium at an initial OD_600_ of 0.02 with or without arabinose to construct SRP^+^ and SRP^−^ cells, respectively. When cultures reached an OD_600_ ~0.4-0.5 (SRP^−^ cells were cultured for 2-3 h), protein expression was induced by 0.5 mM IPTG, and 100 μM biotin was also added at this point. After 3 h of cultivation, cells were harvested by centrifugation. For Ffh depletion, 0.2% glucose was added 2 h before harvesting cells. Then the samples were analyzed by SDS-PAGE and immunoblotting. Biotinylated proteins were detected by streptavidin-HRP and the total amount of protein was detected by anti-FLAG antibody. Detection was performed by the DAB substrate kit (Thermo Scientific, USA).

## RESULTS

### Characterization of an SRP Suppressor

A previous study in our laboratory demonstrated that SRP suppressors were all associated with protein translation (Zhao et al., 2021). To determine whether suppressor mutations of SRP are all mapped to chromosomal loci that influence protein translation and whether there is an alternative pathway to transport SRP substrates when the SRP pathway is blocked, we used the same suppressor approach as previously described to screen SRP suppressors (Zhao et al., 2021). We obtained another suppressor strain MY1901 that could survive when SRP was deleted (**Figure 1A**). The growth rate of MY1901 was significantly reduced compared with that of wild-type strain MG1655, demonstrating that the MY1901 strain had a severe growth defect. We also found a longer lag time during the growth course of MY1901 than that of MG1655 (**Figure 1A**), indicating that the lag time before regrowth bought time for cell adaptation in the absence of SRP. Whole-genome sequencing of the suppressor strain MY1901 and the original strain MG1655 allowed us to identify the suppressor mutation located in the SD sequence of ribosome S10 operon (**Figure 1B** and **Supplementary Table S2**). The S10 operon encodes 11 different ribosomal proteins (Zengel and Lindahl, 1994). To determine whether restoration of the wild-type alleles reverts the MY1901 strain to the wild-type growth phenotype, the Ffh expression and reverting suppressor mutation to the wild-type allele in the MY1901 strain were carried out (**Figure 1C**). The expression of Ffh markedly shortened the lag time and increased the growth rate (**Figure 1D**). Although reverting the suppressor mutation to the wild-type allele further increased the cell growth rate, the growth rate of strain MY1901FS was not equal to that of the wild-type strain MG1655 (**Figure 1D**). We also found that the growth rate of the MY1901 strain carrying the empty vector pTrc99K was a two-fold decrease relative to that of the MY1901 strain without any plasmids (**Figure 1A, D**). Therefore, the plasmid pTrc99K caused a significant burden on cell growth of strain MY1901, which resulted in the growth rate of strain MY1901FS did not fully recover to that of the wild-type strain. Thus, the deletion of Ffh and suppressor mutation indeed reduced cell growth. These results suggested that the mutation in the SD sequence of the S10 operon contributed to the cell growth without SRP and this mutation was a novel suppressor of SRP.

**FIGURE 1.**
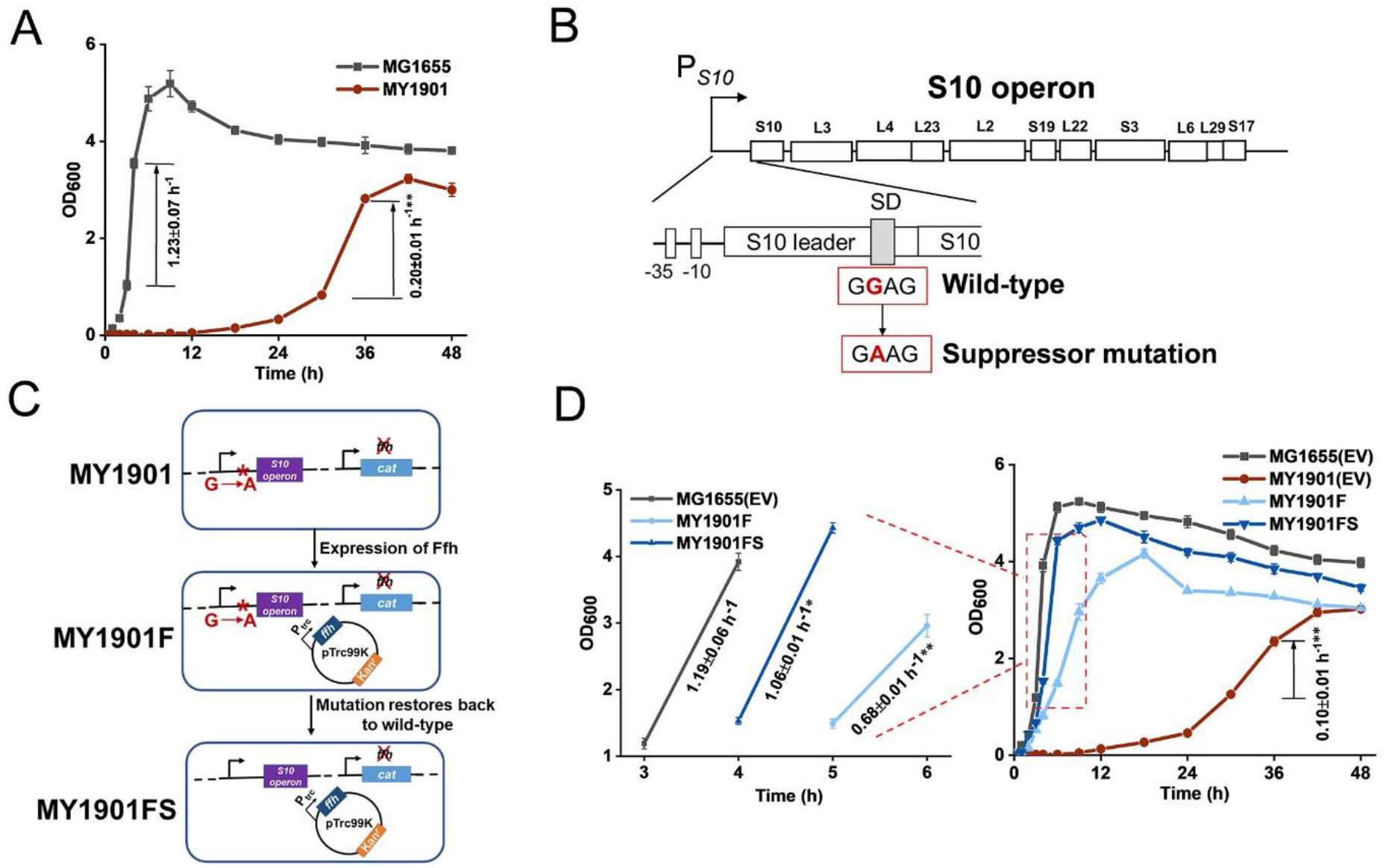
Identification of SRP suppressor strain and its growth. **(A)** Growth curves and growth rates of the wild-type strain MG1655 and the suppressor strain MY1901. **(B)** The suppressor mutation sites in the SD sequence of S10 operon. The region and encoded ribosomal proteins of the S10 operon are shown. The wild-type SD sequence (GGAG) mutated into suppressor mutation (GAAG). **(C)** Restoration of the wild-type allele. Plasmid pTrc99K-Ffh was first transformed into strain MY1901, generating the strain MY1901F. Ffh protein was produced at low level in MY1901 to reduce the toxicity of overexpression of Ffh. Then, the suppressor mutation (GAAG) was restored to wild-type SD sequence (GGAG) by site-directed mutagenesis, generating the strain MY1901FS. **(D)** Growth curves and growth rates of four strains: MG1655 and MY1901carrying the empty vector (EV) pTrc99K, MY1901F, and MY1901FS. Strains MG1655 and MY1901 carrying the empty vector pTrc99K were used as control. Solid curves are the mean of three independent measurements, and error bars represent the standard deviation of the mean value. Growth rates were figured out from the logarithmic phase. All growth rates shown represent the mean ± standard deviation of three independent experiments. Statistical significance compared to wild-type strain using Student’s *t*-test is indicated as follows: *, *P* 0.05; **, *P* 0.01.

We hypothesized that the evolutionarily selective forces shaped the translation process during screening suppressors in SRP-deletion cells. Thus, we tried to improve cell growth through laboratory evolution. Strain MY1091 was evolved through 80 rounds of serial passage in the absence of any other selective pressure. The evolved strain showed an insignificant change in the growth rate but a shorter lag time before regrowth and increased biomass compared with the initial strains (**Supplementary Figure S1**). The laboratory evolved strains were sequenced and new changes in genes were identified (**Supplementary Table S2**). In the evolved strain MY1901, we found a mutation in the ribosomal protein S10 (RpsJ) belonging to the S10 operon, confirming that the S10 operon plays an important role in cell survival. Additionally, we obtained mutations in the DNA replication initiator protein DnaA (Hughes et al., 1988) and RNA polymerase RpoBC operon (Ishihama and Fukuda, 1980) (**Supplementary Table S2**), which regulate gene expression. Taken together, protein synthesis plays a critical role in suppressing the loss of SRP.

### Effects of SRP Suppressor on Ribosomal Protein Biogenesis

To examine the differences between wild-type and mutant SD sequence, the mfold web server (Zuker, 2003) was used to model mRNA structures of the S10 leader sequence containing the SD sequence. S10 leader is required for the regulation of S10 operon (Allen et al., 1999). We found a 2.4 kcal mol^-1^ difference in the minimum free energy of the thermodynamic ensemble of the predicted structure of mutant SD compared to the wild-type SD (**Supplementary Figure S2**), suggesting that the S10 leader with mutant SD was less stable. Because translation initiation is partially modulated by the SD sequence (Yang et al., 2016), the effects of the mutant SD sequence on protein translation were determined. The expression level of the green fluorescent protein (GFP) was used to characterize protein translation levels in cells. Regardless of whether under the original promoter P*_S10_* or the arabinose inducible promoter P*_araBAD_*, the expression level of GFP with the mutated SD (SD*) was significantly decreased (**Figure 2A**), confirming that the suppressor mutation weakened the binding of mRNA SD sequence to rRNA in ribosomes. This result suggested that the translation initiation rate of the S10 operon was reduced. To examine whether the suppressor mutation decreases the abundance of ribosomal proteins, the whole-cell lysates proteome of strain MY1901 was analyzed (**Supplementary Data Set S1A**). The expression level of each gene was normalized by that in wild-type strain MG1655(Zhao et al., 2021). Unexpectedly, the levels of the S10 operon and even the overall ribosomal proteins were upregulated (**Figure 2B** and **Supplementary Data Set S1B**). Given that the growth rate is linearly correlated with the cell’s active ribosome content (Scott et al., 2010) and the suppressor strain MY1901 showed a very low growth rate of 0.2 h^-1^(**Figure 1A**), the active ribosomal protein content would be reduced. We speculated that the increased pool of ribosomal proteins may be caused by the accumulation of ribosomal proteins. There are two possibilities: one is that the ribosomes stalls at the translation initiation site (Zhao et al., 2021); the other is that cellular stress responses caused by the decreased active ribosome level induce the upregulation of ribosomal proteins, including the S10 operon. Thus, further studies are needed for a better understanding of the biogenesis of ribosomal proteins.

**FIGURE 2.**
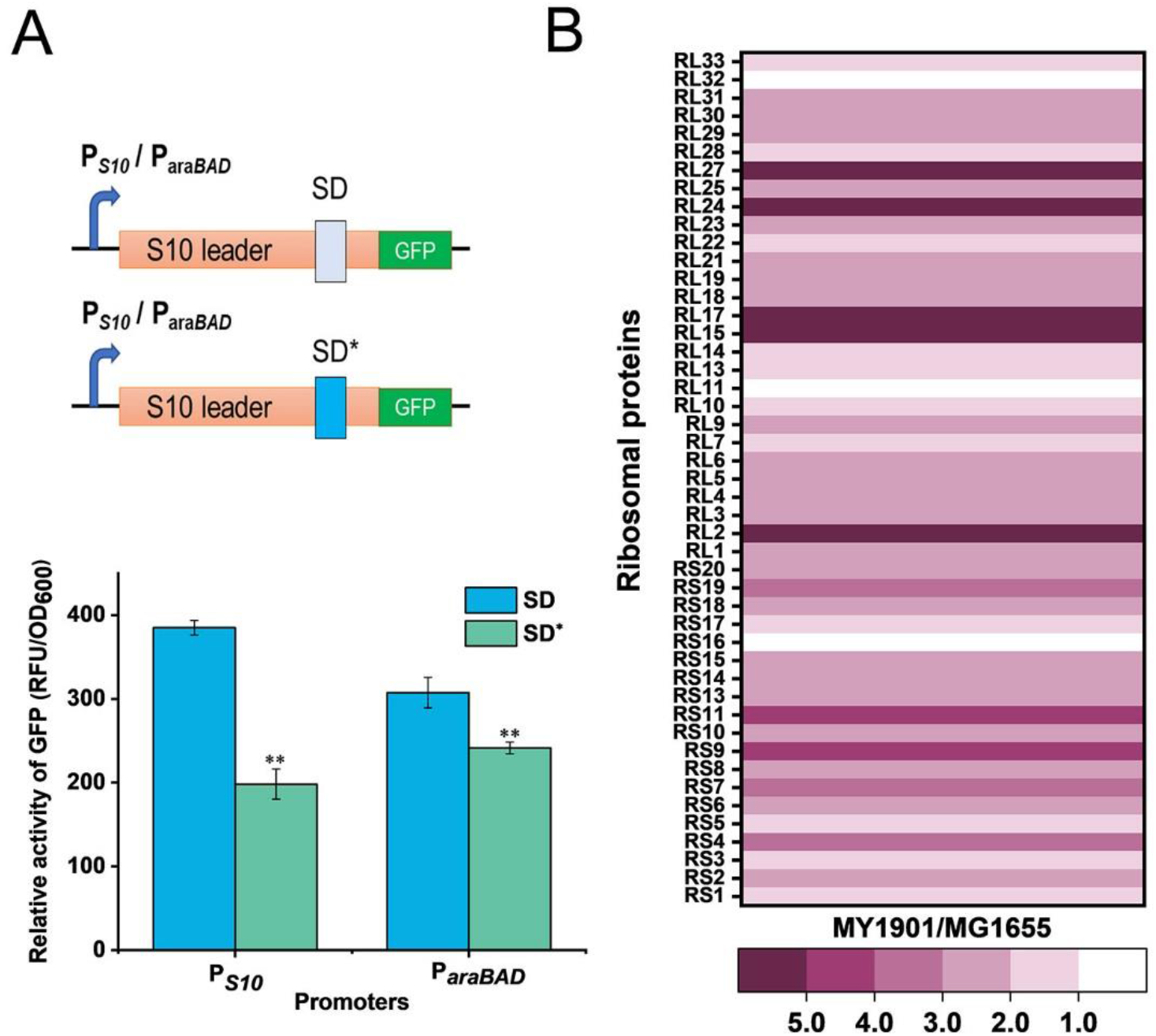
Effects of SRP suppressor on ribosomal protein synthesis. **(A)** Effects of mutant SD sequence on protein translation. GFP was used as a reporter to measure gene expression. (Top) Graphical representation of GFP expression. P*_S10_* / P*_araBAD_*-GFP with wild-type SD and suppressor mutation SD* were introduced into cells. (Bottom) Fluorescence measurement of GFP. Data represent the mean of six independent experiments, and error bars represent the standard deviation of the mean value. Statistical significance compared to wild-type SD sequence using Student’s *t*-test is indicated as follows: *, *P* 0.05; **, *P* 0.01. **(B)** Fold changes in the expression of ribosomal proteins in strain MY1901relative to that in strain MG1655 (**Supplementary Data Set S1B**).

### Translation Speed and Accuracy in Suppressor Cells

In order to test the potential relationship between the biogenesis of ribosomal proteins and protein translation, we first carried out polysome profiling, which could detect various defects of translation. The polysome profile of suppressor cells was changed relative to that of wild-type cells (**Figure 3A**). The 30S to 50S (30S/50S) ratio was not significantly altered in suppressor and wild-type strains, but 30S and 50S peaks of suppressor cells were higher than those of wild-type cells (**Figure 3A**), indicating an increase in the proportion of free subunits. The 70S peak of the wild-type cells was almost indistinguishable from that of suppressor cells (**Figure 3A**), suggesting that some 70S ribosomes paused at the translation initiation site, which can cause the ribosomal proteins accumulated in the cytoplasm (Zhao et al., 2021). We also observed that the polysome to 70S monosome (P/M) ratio was decreased in suppressor cells relative to that in wild-type cells (**Figure 3A**), suggesting that translation elongation was inhibited in suppressor cells. This result also demonstrates that the active ribosomal protein content in suppressor cells is reduced relative to that in wild-type cells. Thus, the protein translation efficiency is affected in suppressor cells.

**FIGURE 3.**
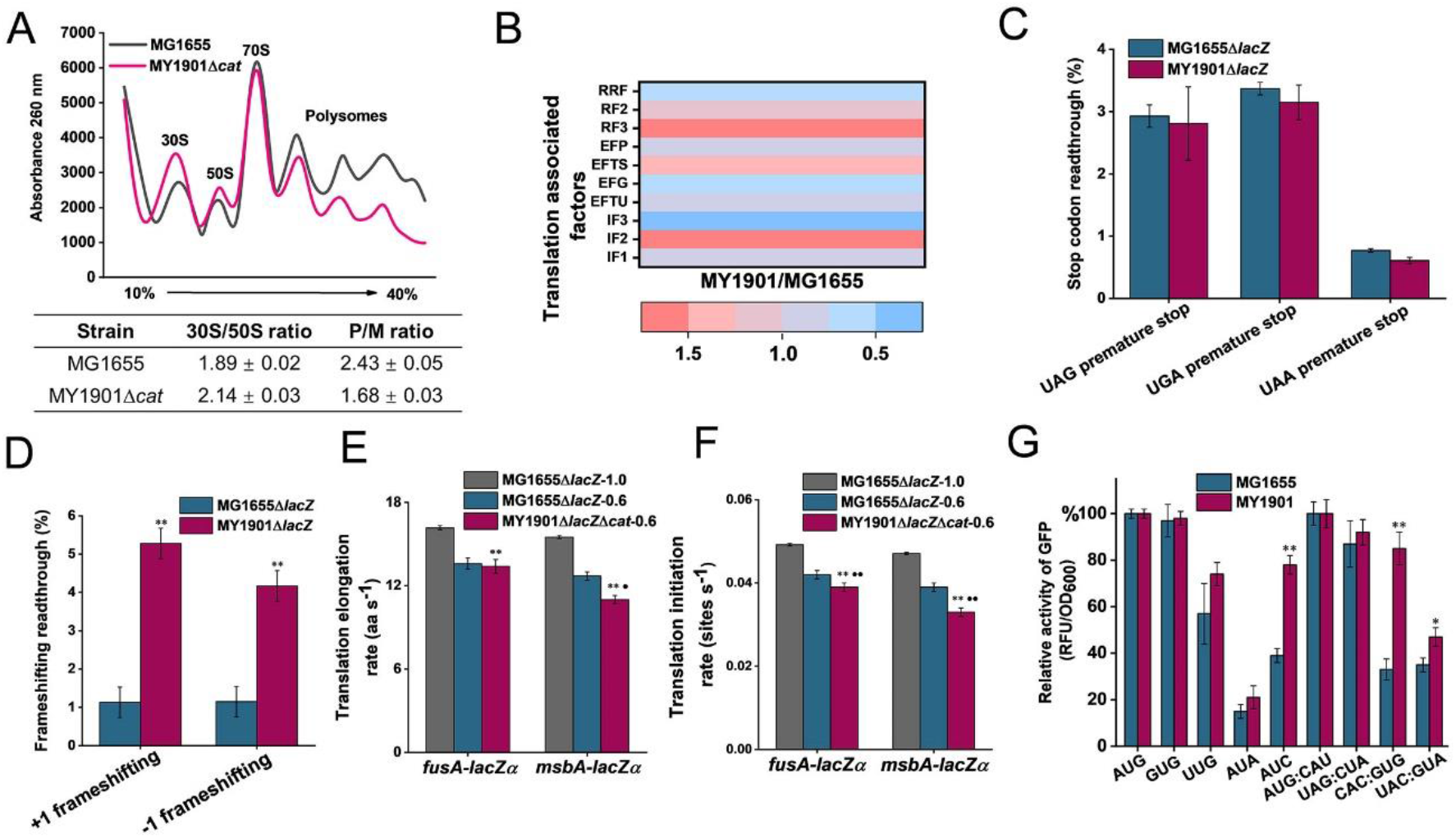
Effects of SRP suppressor on translation efficiency. **(A)** Polysome profiles of the wild-type strain MG1655 and the suppressor strain MY1901Δ*cat*. Data represent the mean ± standard deviation of three independent experiments. **(B)** Fold changes in the expression of translation associated factors in strain MY1901relative to that in MG1655 (**Supplementary Data Set S1B**). **(C-D)** Effect of suppressor mutation on stop-codon **(C)** and frameshifting **(D)** readthrough. All data are averages of at least three independent experiments, and error bars represent the standard deviation of the mean. β-galactosidase activities of strains harboring the mutant LacZ reporters were calculated relative to those of the same strain harboring the wild-type LacZ. Statistical significance compared to the suppressor strain MY1901Δ*cat* using Student’s *t*-test is indicated as follows: *, *P* 0.05; **, *P* 0.01. **(E)**The translation elongation rates of FusA-LacZα and MsbA-LacZα under different growth rates (MG1655Δ*lacZ*, 1.0 h^-1^; MG1655Δ*lacZ*, 0.6 h^-1^; MY1901Δ*lacZ*Δ*cat*, 0.6 h^-1^). **(F)** The translation initiation rate of proteins was estimated based on their corresponding translation elongation rate. All data are averages of three independent experiments, and error bars represent the standard deviation of the mean value. Statistical significance compared to the strain MG1655Δ*lacZ* with growth rate 1.0 h^-1^ using Student’s *t*-test is indicated as follows: *, *P* 0.05; **, *P* 0.01. Statistical significance compared to the strain MG1655Δ*lacZ* with growth rate 0.6 h^-1^ using Student’s *t*-test is indicated as follows: •, *P* 0.05; ••, *P* 0.01. (**G**) Initiation with various non-canonical start codons and non-initiator tRNAs codons in the wild-type and suppressor strains. Relative activities of mutant GFP were calculated relative to that of the same strain harboring the wild-type GFP. The wild-type start codon was indicated as AUG, and the wild-type initiator tRNA was indicated as AUG: CAU. All data are averages of three independent experiments, and error bars represent the standard deviation of the mean value. Statistical significance compared to the strain MG1655 using Student’s *t*-test is indicated as follows: *, *P* 0.05; **, *P* 0.01.

Translation can be divided into four phases: initiation, elongation, termination, and ribosome recycling (Rodnina, 2018). We next compared the abundances of translation-associated factors via whole-cell lysates proteome analysis. Ribosome recycling factor (RRF) and elongation factor-G (EF-G) are involved in ribosome recycling (Prabhakar et al., 2017). The expression of RRF and EF-G was downregulated in the suppressor strain (**Figure 3B** and **Supplementary Data Set S1B**), indicating that the ribosome recycling was reduced, which may be caused by decreasing translation in suppressor cells. Although the expression of release factors mediating translation termination was upregulated, the efficiency of translation termination at three stop codons was only slightly increased (**Figure 3B** and **C**). This data showed that the translation termination rate was unaffected in suppressor cells. We observed that the protein abundance of elongation factors in suppressor cells was inconsistent and changed little in comparison with that in wild-type cells (**Figure 3B** and **Supplementary Data Set S1B**). To examine the efficiency of translation elongation, we measured its fidelity and speed through GFP fluorescence intensity measurement and LacZα induction assay, respectively. In strain MY1901, we observed a marked increase in translational frameshifting readthrough relative to that in the wild-type strain (**Figure 3D**), suggesting that the fidelity of translation elongation was decreased. Additionally, a LacZ induction assay was used to measure the translation elongation speed (Zhu et al., 2016). We used a cytoplasmic protein FusA and an inner membrane protein MsbA to define the level of translation elongation rate (**Supplementary Figure S3**). When cells were grown at the same rich growth media (Glucose + cAA), the growth rates of wild-type and suppressor cells were approximately 1.0 h^-1^ (Zhao et al., 2021) and 0.6 h^-1^, respectively (**Supplementary Figure S3A** and **Table S3**). As expected, the translation elongation rate of suppressor cells was markedly decreased compared with that of wild-type cells (**Figure 3E** and **Supplementary Figure S3B** to **F**). As translation elongation rate closely depends on growth rate (Dai et al., 2016), we further reduced the growth rate of wild-type cells to 0.6 h^-1^ that was similar to that of suppressor cells (**Supplementary Figure S3A**). The elongation rate of the MsbA in suppressor cell was reduced by approximate 2.0 aa s^-1^ (amino acids per second) compared with that in wild-type cells, but the elongation rate of the FusA was similar in both wild-type and suppressor cells (**Figure 3E** and **Supplementary Figure S3B** to **F**). We also observed that the elongation rate of MsbA was slower than that of FusA (**Figure 3E** and **Supplementary Figure S3B** to **F**), which is consistent with the observation that the translation elongation speed of inner membrane proteins is slowed down during targeting, but not that of cytoplasmic proteins (Fluman et al., 2014). These data suggested that the effect of selectively reducing translation elongation rate of inner membrane proteins was enhanced in suppressor cells.

For the expression of translation initiation factors in suppressor cells, IF1 and IF3 were downregulated, but IF2 was upregulated (**Figure 3B**). Given that IF2 accelerates ribosomal subunit joining, whereas IF1 and IF3 slowed down subunit association (Ling and Ermolenko, 2015; Naaktgeboren et al., 1977), the capacity of formation of 70S initiation complex may not be significantly impaired, although cells showed severe defects in growth. We next examined the translation initiation rate by a computational model called the homogeneous ribosome flow model (HRMF), in which the translation elongation rate is assumed to be constant (Margaliot and Tuller, 2012). The translation initiation rate can be estimated by the measurable translation rate and translation elongation rate (**Supplementary Table S4**). The translation initiation rate in suppressor cells showed a similar trend to the translation elongation rate. The initiation rate of FusA was not markedly changed in the suppressor and wild-type cells when grown at the same growth rate (**Figure 3F**). However, the inner membrane protein MsbA had a slower translation initiation rate in suppressor cells than that in wild-type cells when grown at the same growth rate (**Figure 3F**). These results suggested that the translation initiation rate was selectively decreased in suppressor cells. Thus, in suppressor cells, the translation initiation process was negatively affected, although the formation of the 70S initiation complex was not markedly influenced (**Figure 3A**). As translation initiation factors play a vital role in translation initiation fidelity (Ling and Ermolenko, 2015), we addressed whether the suppressor was detrimental to the fidelity of start codon selection and initiator tRNA binding. We changed the start codon of GFP from AUG to other two canonical start codons GUG and UUG, and two near cognates AUA and AUC (Hecht et al., 2017); and the initiator tRNA codon of GFP from AUG: CAU to non-initiator tRNAs UAG: CUA, CAC: GUG and UAC: GUA. We measured the GFP fluorescence of these GFP variants in the wild-type and suppressor cells. We observed that the expression levels of GFP with three canonical start codons (AUG, GUG, UUG) and a near cognate (AUA) were similar in both wild-type and suppressor strains (**Figure 3G**). However, the expression level of GFP with the near cognate AUC as the start codon in the suppressor strain was significantly increased relative to that in the wild-type strain (**Figure 3G**). We also found that in the suppressor strain, the expression levels of GFP with non-initiator tRNAs CAC: GUG and UAC: GUA were significantly increased compared with those in the wild-type strain (**Figure 3G**). Additionally, the level of GFP with the initiator tRNA AUG: CAU in the suppressor strain was not changed relative to that in the wild-type strain (**Figure 3G**). Thus, the fidelity of translation initiation in the suppressor cells was decreased compared with that in wild-type cells. Taken together, the suppressor cell trades translation speed and accuracy for cell survival in the absence of SRP.

### SRP-dependent Protein Targeting in Suppressor Cells

In *E. coli*, most inner membrane proteins are delivered by the co-translational SRP pathway (Elvekrog and Walter, 2015). In principle, SRP-dependent proteins are not properly targeted to the bacterial cytoplasmic membrane after Ffh depletion (Bernstein and Hyndman, 2001; Wickström et al., 2011). To test whether the suppressor mutation could suppress protein targeting defects, we first examined cell morphological changes by scanning electron microscopy (SEM) and transmission electron microscopy (TEM). SEM images showed that the suppressor strain MY1901 still had typical rod morphology but had a rougher surface relative to the wild-type strain MG1655 (**Figure 4A** and **Supplementary Figure S4**). TEM images showed that the suppressor strain MY1901 retained cell wall integrity but had damaged inner membrane structure (**Figure 4A** and **Supplementary Figure S4**). MY1901 displayed a significant detachment of the inner membrane from the outer membrane (**Figure 4A** and **Supplementary Figure S4**). Thus, the suppressor mutation partially offsets the negative effects of the loss of the SRP pathway on the inner membrane protein translocation.

**FIGURE 4.**
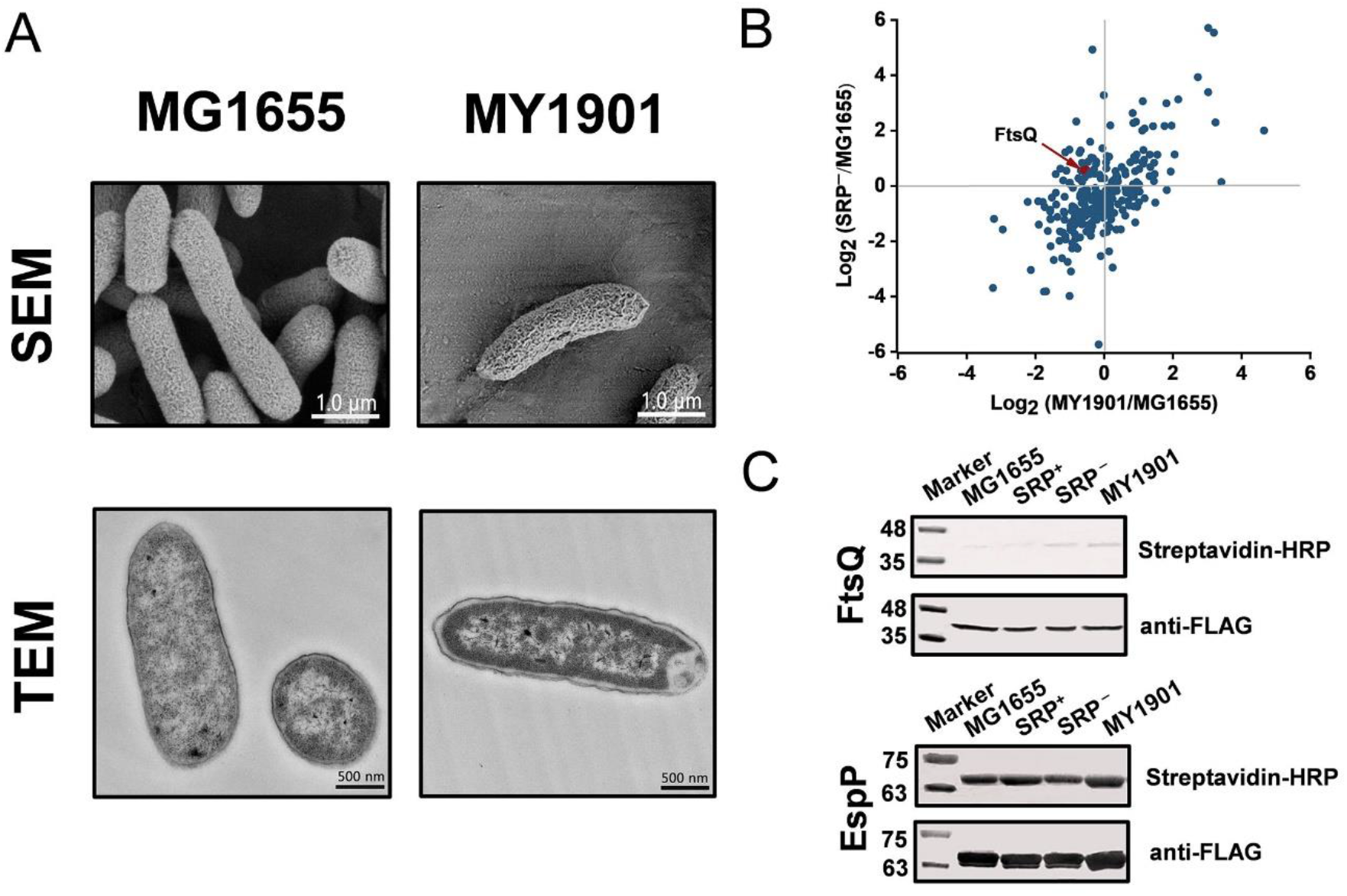
Suppressor mutation suppresses targeting defects of partial inner membrane proteins. **(A)** Scanning electron microscopy (SEM) and transmission electron microscopy (TEM) analysis of the wild-type strain MG1655 and the suppressor strain MY1901. For SEM, the scale bar is 1.0 μm. For TEM, the scale bar is 500 nM. **(B)** Quantification of identified SRP-dependent inner membrane proteins in strains MY1901 and SRP^−^ (**Supplementary Data Set S1D**). Protein abundance of MY1901 and SRP^−^ is relative to that of wild-type MG1655. **(C)** FtsQ (left) and EspP (right) targeting assay by their biotinylation. SRP^+^, Ffh expression in HDB51 strain; SRP^−^, Ffh depletion in HDB51 strain.

To gain an insight into the localization of inner membrane proteins, we performed proteomic analysis of inner membrane proteins in the wild-type strain MG1655 (Zhao et al., 2021) and the suppressor strain MY1901 (**Supplementary Data Set S1C**). The *E. coli* HDB51 was used as a control strain, in which the expression of Ffh was induced by arabinose (Lee and Bernstein, 2001). Depleted Ffh can be obtained after several hours of incubation in the presence of glucose, thus yielding the SRP^−^ strain (Zhang et al., 2012). The inner membrane proteome analysis of SRP^−^ was also conducted (**Supplementary Data Set S1C**). According to our previous study, we identified 262 SRP-dependent inner membrane proteins (Zhao et al., 2021). Our previous study has shown that the inner membrane proteins with a high abundance, such as proteins C_4_-dicarboxylate sensor kinase DcuS and zinc transporter FieF, can be localized to the membrane (Zhao et al., 2021). This suggested that the high protein abundance can be used as an indicator of protein localization. We found that the abundance of many identified SRP-dependent inner membrane proteins in both MY1901 and SRP^−^ cells were higher than their abundance in wild-type cells (**Figure 4B** and **Supplementary Data Set S1D**), indicating that these inner membrane proteins can target to the cytoplasmic membrane in the absence of SRP. This result is consistent with previous studies showing that inhibition of the SRP pathway only partially impedes inner membrane protein targeting (Bernstein and Hyndman, 2001; Newitt et al., 1999; Ulbrandt et al., 1997b). We also found that more proteins were successfully targeted in the MY1901strain than in the SRP^−^ strain (**Figure 4B**), suggesting that the suppressor mutation indeed plays a role in inner membrane protein targeting without SRP. FtsQ is an SRP-dependent protein, which is often used as a model protein for studying SRP-mediated protein targeting (Scotti et al., 1999; Zhang and Shan, 2012). However, the targeting of FtsQ was not significantly inhibited in strains SRP^−^ and MY1901 (**Figure 4B** and **Supplementary Data Set S1D**). To examine the targeting level of FtsQ, we used a sensitive method based on protein biotinylation (Jander et al., 1996; Zhang and Shan, 2012). A small biotinylatable peptide Avi-tag was fused to the periplasmic domain of the targeted proteins. The biotinylated proteins would be the untargeted proteins in which the periplasmic domains are exposed in the cytosol. Thus, the protein biotinylation can be used for protein targeting assay. However, in contrast to the prediction of proteomic analysis, the FtsQ targeting showed a slight defect in both the SRP^−^ and MY1901 strains (**Figure 4C**), suggesting that the suppressor mutation played little role in the targeting of FtsQ. We also found that the targeting levels of the SRP-independent protein EspP were similar in both wild-type and MY1901strains (**Figure 4C**), suggesting that the targeting of SRP-independent proteins was not affected by the loss of SRP. However, there was a slight defect in EspP targeting in SRP^−^ strain. This may be caused by the secondary effect due to the defects of SRP-dependent transporters, such as SecY, SecG, and YajC (**Supplementary Figure S5A** and **Data Set S1E**). We also found that the protein abundance of almost all identified membrane components of transporters in MY1901 was not lower than that in the SRP^−^ strain (**Supplementary Figure S5A** and **Data Set S1E**). These results indicated that the SRP suppressor partially contributed to inner membrane protein targeting and allowed for targeting of some SRP-dependent proteins without causing a failure of targeting of SRP-independent proteins.

Additionally, the expression of heat shock response related chaperones and proteases was not upregulated (**Supplementary Figure S5B** and **Data Set S1B**), suggesting that the heat shock response played little role in compensating the loss of SRP, which is consistent with our previous study (Zhao et al., 2021). In strain MY1901, the protein abundance of SecA was not affected and other transport components such as SecYEG, YajC, SecD, YidC, SecF, and FtsY showed a decreased level relative to that in the wild-type strain (**Supplementary Figure S5A** and **Data Set S1E**). Moreover, we found that the protein abundance of SecF in the MY1901 strain was significantly decreased relative to that in the SRP^−^ strain (**Supplementary Figure S5A** and **Data Set S1E**). This suggested that the component of the Sec translocon SecF may not be involved in the protein targeting process without SRP. In contrast, the protein abundance of SecY and FtsY in MY1901 was two times higher than that in the SRP^−^ strain, which is likely caused by the effective targeting of inner membrane proteins with the assistant of translational control. Overall, protein transport components were unlikely to play a major role in mediating SRP-dependent protein targeting in the absence of SRP.

## DISCUSSION

Co-translational protein targeting by SRP is an essential and conserved pathway that delivers most inner membrane proteins to their correct subcellular destinations (Saraogi and Shan, 2014). Our previous work revealed that SRP was not essential in *E. coli* when the translation initiation and elongation rate were decreased (Zhao et al., 2021). Isolation of suppressors is a useful strategy to provide insight into certain molecular mechanisms by suggesting which cellular component is involved in an inefficient process (Lee and Beckwith, 1986). The SRP suppressors involved in protein translation initiation have been identified before, and these suppressors affect the translation process (Zhao et al., 2021). In this study, we obtained an SRP suppressor associated with protein translation too. The regulation of translation may be a general way to mediate the translocation of SRP-dependent proteins in the absence of SRP.

We observed that in suppressor cells, the ribosomal protein expression was upregulated (**Figure 2B**) and the 30S and 50S ribosomal subunits accumulated (**Figure 3A**), but the content 70S ribosome complex was not markedly changed relative to those in the wild-type strain (**Figure 3A**). This led us to propose that the increased ribosomes are inactive and accumulate in the cytosol. Furthermore, in earlier works, deletion of SRP caused the downregulation of ribosomal proteins (Wickström et al., 2011; Zhang et al., 2012), which suggested that the absence of SRP alone cannot increase the level of ribosomal proteins. Thus, the SRP suppressor and cellular stress responses may play an important role in ribosomal protein synthesis.

In suppressor cells, the protein translation initiation was impeded (**Figure 3F**), but the initiation time of translation was constant in wild-type and suppressor cells under different growth rates (**Supplementary Table S3**), which suggested that the pausing at the start of the initiation can be negligible, and the process of 70S ribosome complex entry into the elongation cycle is slower in suppressor cells. Thus, the SRP suppressor may be associated with the transition from initiation to elongation. The closely related relationship between translation initiation and elongation (Riba et al., 2019) and the decreased translation initiation and elongation rates caused by the absence of SRP (**Figure 3E** and **F**) suggested the possibility that this SRP suppressor mutation can either reduce the translation initiation rate or the elongation rate.

Furthermore, we showed that the translation fidelity was decreased in suppressor cells (**Figure 3D** and **G**). Because the fidelity of translation initiation is modulated by the initiation factors (Ayyub et al., 2017; Caban et al., 2017) and the suppressor mutation was associated with the biogenesis of ribosomal proteins (**Figure 2B**), we speculated that the suppressor may indirectly regulate the fidelity of translation initiation by influencing the abundance of translation initiation factors (**Figure 3B**). We observed that the fidelity of translation elongation was also decreased, implying that suppressor mutation may inactivate the quality control system. Earlier works revealed that mistranslation could provide a growth advantage in response to stress (Gu et al., 2014; Mohler and Ibba, 2017). Hence, the decreased fidelity of translation initiation and elongation may result from the SRP deletion stress response.

Increasing evidence has supported the notion that the translation elongation of nascent polypeptide regulates the targeting of SRP-dependent proteins (du Plessis et al., 2011; Zhang and Shan, 2012), thus decreasing the elongation rate contributes to the survival of SRP deletion cells. Decreasing the translation elongation rate extends the time window for protein targeting, which plays a critical role in suppressing the loss of SRP (Zhao et al., 2021). SRP binds to the ribosome-nascent chain complex when the N-terminus of the first TMD is exposed from the ribosome. The maximal SRP binding site is 55 amino acids from the ribosomal peptidyl transferase center in *E. coli* (Schibich et al., 2016). Assuming that ~30 amino acids can fit into the ribosome exit tunnel (Bornemann et al., 2008), 25 residues would be exposed outside the tunnel. At a translation elongation rate of ~15 aa s^-1^ in rapid growth conditions (Zhao et al., 2021), the maximum time required for protein localization is ~2 s (**Figure 5**). Thus, with the help of SRP, most translating ribosomes move to the membrane within this period in *E. coli*. Without SRP, suppressors slowed the translation elongation rate to ~11 aa s^-1^(**Figure 3E** and **Supplementary Table S3**), which provides ~2 s for nascent chains of 55 amino acids to target to the inner membrane (**Figure 5**). However, it is not likely that nascent chains successfully target to the membrane within ~2 s without SRP. To get a longer time to find the membrane, the length of translating nascent chains is more likely longer than 55 amino acids. However, the nascent chain cannot exceed a specific length as aggregation would prevent protein from being targeted (Flanagan et al., 2003; Siegel and Walter, 1998), and this specific length is called the critical length (L) for targeting. Proteins with fewer transmembrane domains (TMDs) or longer first loop lengths have a longer critical length (Zhao et al., 2021). Furthermore, if the nascent chain exceeds a critical length of ~140 amino acids, it becomes translocation-incompetent (Flanagan et al., 2003; Siegel and Walter, 1998). In suppressor cells, the upper limit of the critical time for protein targeting would be ~10 s (**Figure 5**). If the targeting time of some SRP-dependent proteins exceeds 10 s, these proteins would not be targeted to the inner membrane in suppressor cells. Thus, the suppressor extends the time window to ~2-10 s **(Figure 5**). Taken together, this model shows that SRP greatly shortened the protein targeting time by 8 s, which minimizes the cost of targeting and maintains fast growth. Overall, our data suggest that in response to the deletion of SRP, suppressor cells attenuate translation elongation to give the translating ribosomes more time to find and target to the inner membrane.

**FIG 5.**
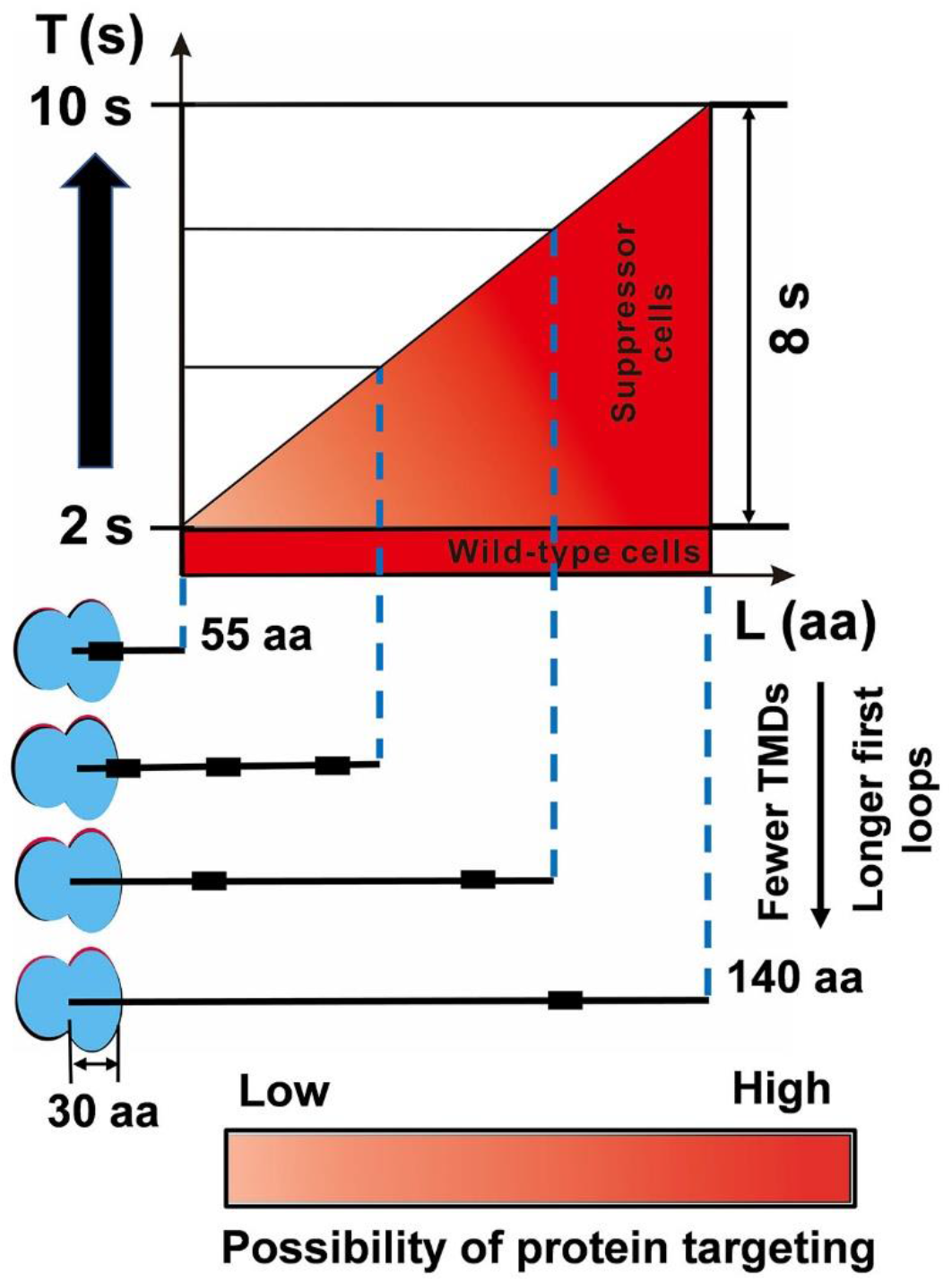
The SRP suppressor extends the time window for protein targeting. In wild-type MG1655 cells, although the critical length (L) of inner membrane proteins is variable, the targeting time (T) is fast (~2 s). The suppressor cells with a slower elongation rate extended the time window for protein targeting to 2-10 s. For details and references, see Discussion.

As expected, the suppressor mutation can partially offset the defective targeting of inner membrane proteins (**Figure 4B**), which is consistent with the previous result (Zhao et al., 2021). Furthermore, we found that in the SRP depletion strain SRP^−^, many proteins can successfully target to the inner membrane (**Figure 4B**). However, proper localization of these proteins cannot bypass the requirement of SRP (Phillips and Silhavy, 1992). We speculated that the proteins that could be correctly located in the suppressor strain MY1901 but not in SRP depletion strain SRP^−^ may be responsible for cell survival. We hypothesized that specific membrane protein targeting defects could block the essential cellular process, which would be responsible for the loss of cell viability. Among these localization defective proteins, only one protein PgsA is essential for *E. coli* (**Supplementary Data Set S1F**). PgsA catalyzes the step in the synthesis of the acidic phospholipids that are considered to be indispensable in multiple cellular processes (Gopalakrishnan et al., 1986; Kikuchi et al., 2000). We inferred that mislocalization of PgsA inhibited cell growth. Furthermore, in SRP^−^ strain, some transportation associated proteins (AsmA, Bcr, PheP, YbaL, and YidE) did not successfully target to the inner membrane (**Supplementary Data Set S1F**), which would impair membrane traffic and decrease energy production. More studies are needed to investigate the targeting of some proteins that determine whether cells can survive without SRP.

## SUMMARY

The signal recognition particle (SRP)-dependent delivery pathway is essential for membrane protein biogenesis. Previously, we reported that SRP was nonessential in *Escherichia coli*, and slowing translation speed played a critical role in membrane protein targeting. Here we identified a novel SRP suppressor that is also involved in translation. We found that translation speed and accuracy regulate membrane protein targeting. A slowdown of translation speed extended the time window for protein targeting. Meanwhile, a moderate decrease in translation fidelity ensured a suitable translation speed for better cell growth. These results argued that translation control could be a practical way to compensate for the loss of SRP.

## DATA AVAILABILITY

The datasets generated and analyzed during the current study are included in this published article.

## AUTHOR CONTRIBUTION

LZ, GF and DZ designed the experiments. LZ and YC performed experiments. LZ, YC, ZX, and TC analyzed the data. LZ and DZ wrote the manuscript

## ACKNOWLEDGMENTS

We thank Dr. Manlu Zhu (Central China Normal University) for plasmid pKUT15-fusA-lacZα, Dr. Changhao Bi (TIB, CAS) for plasmid pRed_Cas9_ΔpoxB300, Dr. Harris D Bernstein (NIH) for strain HDB51 and plasmid pJH29 and Dr. Zhidan Zhang (TIB, CAS) for help with mass spectrometry.

