## Supplementary material for "Compensating complete loss of signal recognition particle during co-translational protein targeting by the translation speed and accuracy": Supplememtary Table S1

### Supplementary Figures and Tables

#### Supplementary Table S1A. Strains used in this study.

| Strain | Genotype | Source or Reference |
|---|---|---|
| MG1655 | K-12 $F^-$ $\lambda^-$ rph-1 | (Blattner et al., 1997) |
| HDB51 | WAM113 *secB*$^+$ *zic-4901*::Tn10 | (Lee and Bernstein, 2001) |
| MY1410 | MG1655 *ffh*::Cm$^r$, pKDFB | (Zhao et al., 2021) |
| MY1901 | MG1655 *ffh*:: Cm$^r$, C3453283T | This study |
| MY1901F | MY1901, pTrc99K-Ffh | This study |
| MY1901FS | MY1901F, T3453283C | This study |
| MY1901Δ*cat* | MY1901, Δ*cat* | This study |
| MG1655Δ*lacZ* | MG1655, Δ*lacZ* | (Zhao et al., 2021) |
| MY1901Δ*lacZ*Δ*cat* | MY1901, Δ*lacZ*Δ*cat* | This study |

#### Supplementary Table S1B. Plasmids used in this study.

| Plasmid | Relevant genotype | Source or Reference |
|---|---|---|
| pTrc99K-Ffh | pTrc99K, $P_{trc}$-*ffh*, Km$^r$ | (Zhao et al., 2021) |
| pKUT15-fusA-lacZα | OriSC101 lacIq $P_{lac}$-*fusA-lacZα*, $P_{tet}$-lacZω, Gm$^r$ | (Dai et al., 2018) |
| pKUT15-lacZα | pKUT15, $P_{lac}$-*lacZα*, $P_{tet}$-*lacZω* | (Zhao et al., 2021) |
| pKUT15-msbA-lacZα | pKUT15, $P_{lac}$-*msbA-lacZα, $P_{tet}$-lacZω* | (Zhao et al., 2021) |
| pLacZ | pTrc99K, $P_{trc}$-*lacZ* | This study |
| $pLacZ_{835UAA}$ | pLacZ, UAA mutant | (O'Connor et al., 1992) |
| $pLacZ_{12\text{-}6UAG}$ | pLacZ, UAG mutant | (O'Connor et al., 1992) |
| $pLacZ_{34\text{-}11UGA}$ | pLacZ, UGA mutant | (O'Connor et al., 1992) |
| $pLacZ_{lac10(-)}$ | pLacZ, -1 frameshift | (O'Connor et al., 1992) |
| $placZ_{lac7(+)}$ | pLacZ, +1 frameshift | (O'Connor et al., 1992) |
| pRed_Cas9_ΔpoxB300 | OrirepA101, exo, bet, gam, arabinose operon, Cas9, gRNA, Ap$^r$ | (Zhao et al., 2016) |
| pGFP | pTrc99K, $P_{trc}$-$gfp_{AUG}$ | This study |
| $pGFP_{GUG}$ | pTrc99K, $P_{trc}$-$gfp_{GUG}$ | This study |
| $pGFP_{UUG}$ | pTrc99K, $P_{trc}$-$gfp_{UUG}$ | This study |
| $pGFP_{AUA}$ | pTrc99K, $P_{trc}$-$gfp_{AUA}$ | This study |
| $pGFP_{AUU}$ | pTrc99K, $P_{trc}$-$gfp_{AUU}$ | This study |
| $pGFP_{AUG}metY_{CAU}$ | pGFP, expressing $tRNA^{fMet}$ with CAU anticodon | This study |
| $pGFP_{UAG}metY_{CUA}$ | $pGFP_{AUG}metY_{CAU}$, AUG initiation codon mutated to UAG and anticodon of metY was changed from CAU to CUA | This study |
| $pGFP_{CAC}metY_{GUG}$ | $pGFP_{AUG}metY_{CAU}$, AUG initiation codon mutated to CAC and anticodon of metY was changed from CAU to GUG | This study |
| $pGFP_{UAC}metY_{GUA}$ | $pGFP_{AUG}metY_{CAU}$, AUG initiation codon mutated to UAC and anticodon of metY was changed from CAU to | This study |

| | | |
|---|---|---|
| | GUA | |
| Cas9-S10 SD | pRed_Cas9_ΔpoxB300, homologous arms of SD of S10 | This study |
| Cas9-lacZ | pRed_Cas9_ΔpoxB300, homologous arms of lacZ | (Zhao et al., 2021) |
| Cas9-cat | pRed_Cas9_ΔpoxB300, homologous arms of cat | (Zhao et al., 2021) |
| p15A-birA | Orip15A, lacIq $P_{tac}$-*birA*, $Gm^r$ | This study |
| pJH29-EspP-PSBT | pJH29, $P_{tac}$-*espP*, PSBT-FLAG, $Cm^r$ | (Zhang and Shan, 2012) |
| pJH29-EspP-Avi | pJH29, $P_{tac}$-*espP*, Avi-FLAG, $Cm^r$ | This study |
| pJH29-FtsQ-Avi | pJH29, $P_{tac}$-*ftsQ*, Avi-FLAG, $Cm^r$ | This study |
| pJH29-LacZ-Avi | pJH29, $P_{tac}$-*lacZ*, Avi-FLAG, $Cm^r$ | This study |
| pJH30-EspP-PSBT | pJH29, *lac* operon replaced with *araC* operon, $Ap^r$ | (Zhao et al., 2021) |
| pJH30-SD-GFP | pJH30, $P_{araBAD}$-GFP, wild-type SD, $Ap^r$ | This study |
| pJH30-SD*-GFP | pJH30, $P_{araBAD}$-GFP, mutant SD, $Ap^r$ | This study |
| pJH31-SD-GFP | pJH30, $P_{araBAD}$ replaced with $P_{S10,}$ wild-type SD $Ap^r$ | This study |
| pJH31-SD*-GFP | pJH31, $P_{S10}$-GFP, mutant SD, $Ap^r$ | This study |

### Supplementary Table S1C. Primers used in this study.

| Primer | Sequence (5'-> 3') | Use |
|---|---|---|
| pTrc99K-F | GATCCTCTAGAGTCGACCT | Cloning of the vector bone of plasmid pTrc99K |
| pTrc99K-R | GAGCTCGAATTCCATGGTCT | |
| GFP-F | ACAGACCATGGAATTCGAGCTCATGAGTA AAGGAGAAGAACTTTTC | Cloning of *gfp* segment |
| GFP-R | TGCAGGTCGACTCTAGAGGATCCTATTTG TATAGTTCATCCATG | |
| $GFP_{GUG}$-F | ACAGACCATGGAATTCGAGCTCGTGAGT AAAGGAGAAGAACTTTTC | Cloning of $gfp_{GUG}$ with segment |
| $GFP_{UUG}$-F | ACAGACCATGGAATTCGAGCTCTTGAGT AAAGGAGAAGAACTTTTC | Cloning of $gfp_{UUG}$ with segment |
| $GFP_{AUA}$-F | ACAGACCATGGAATTCGAGCTCATAAGT AAAGGAGAAGAACTTTTC | Cloning of $gfp_{AUA}$ with segment |
| $GFP_{AUU}$-F | ACAGACCATGGAATTCGAGCTCATTAGT AAAGGAGAAGAACTTTTC | Cloning of $gfp_{AUU}$ with segment |
| MetY-V-R | GTGAAAACCTCTGACACATG | Cloning of the vector bone of plasmid pGFP with primer pTrc99K-F |
| $MetY_{CAU}$-F | CTGCATGTGTCAGAGGTTTTCACTGGAT CACTATAATGCCTGC | Construction of plamid $pGFP_{AUG}metY_{CAU}$ |
| $MetY_{CAU}$-R | CTATCATGCCATACCGCGAAAGGAGCC CAGTTATTCTGTAGTC | |
| MetY-V-F | CCTTTCGCGGTATGGCATG | |
| $MetY_{CUA}$-F | GTCGGGCTCTAAACCCGAAGATCGT | Construction of plamid $pGFP_{UAG}metY_{CUA}$ |
| $MetY_{CUA}$-R | ACGATCTTCGGGTTTAGAGCCCGACG | |
| $GFP_{UAG}$-F | ACAGACCATGGAATTCGAGCTCTAGAGTAA AGGAGAAGAACTTTTC | |
| $MetY_{GUG}$-F | GTCGGGCTGTGAACCCGAAGATCGT | Construction of plamid $pGFP_{CAC}metY_{GUG}$ |
| $MetY_{GUG}$-R | ACGATCTTCGGGTTCAGAGCCCGACG | |
| $GFP_{CAC}$-F | ACAGACCATGGAATTCGAGCTCCACAGTAA AGGAGAAGAACTTTTC | |
| $MetY_{GUA}$-F | GTCGGGCTGTAAACCCGAAGATCGT | Construction of plamid $pGFP_{UAC}metY_{GUA}$ |
| $MetY_{GUA}$-R | ACGATCTTCGGGTTTAGAGCCCGACG | |
| $GFP_{UAC}$-F | ACAGACCATGGAATTCGAGCTCTACAGTAA AGGAGAAGAACTTTTC | |
| V1-cas9-F | TGAATGGAAGCTTGGATTCTC | Cloning of the vector bone 1 of plasmid Cas9-S10 SD |
| V1-S10-R | TATCCGCCTGAAAGCGTTTGGCTAAGATCTG ACTCCATAAC | |

| | | |
|---|---|---|
| V2-S10-F | CAAACGCTTTCAGGCGGATAGTTTAGAGCTAGAAATAGCAAG | Cloning of the vector bone 2 of plasmid Cas9-S10 SD |
| V2-cas9-R | ACAGGCCCATGGATTCTTCG | |
| S10-UP-F | CGAAGAATCCATGGGCCTGTTTAACCCAGGCTGATCTGC | Cloning of the upstream of the mutated site |
| S10-UP-R | CTGGTCTCATGCAGAACCAAAGAATACGTATCC | |
| S10-DN-F | TGGTTCTGCATGAGACCAGAGCTCCAATTAT | Cloning of the downstream of the mutated site |
| S10-DN-R | AGAATCCAAGCTTCCATTCAAAGAGAAAGCCGGTTTAAGAG | |
| p15-V-F | CCCAGGCATCAAATAAAACG | Cloning of the vector bone of plasmid p15A-birA |
| p15-V-R | GATCCGAATTCCTGCAGTTG | |
| birA-F | TAACAACTGCAGGAATTCGGATCATGAAGGATAACACCGTGCC | Cloning of the *birA* segment |
| birA-R | TTTCGTTTTATTTGATGCCTGGGTTATTTTTCTGCACTACGCAG | |
| 29-V-F | TTCCGGCTTGAACGACATCTTCGAGGCCCAGAAGATCGAGTGGCACGAGGGTTCTGGTGACTACAAAGACGATGACGAC | Cloning of the vector bone of plasmid pJH29-EspP-Avi from pJH29-EspP-PSBT |
| 29-V-R | ATTAGTCCGCCAGTTCCAC | |
| espP-F | TATGTGGAACTGGCGGACTAATATGAATAAAATATACTCTC | Cloning of the *espP* segment |
| espP-R | CGAAGATGTCGTTCAAGCCGGAACCGATATAGTCAGCAGTATTATTC | |
| FtsQ-Avi-V-F | CTTCGAGGCCCAGAAGATCGAGTGGCACGAGGGTTCTGGTGACTACAAAGACGATGACGAC | Cloning of the vector bone of plasmid pJH29-FtsQ-Avi from pJH29-EspP-Avi with primer 29-V-R |
| FtsQ-Avi-F | TATGTGGAACTGGCGGActaatATGTCGCAGGCTGCTCTG | Cloning of the *ftsQ-Avi* tag segment |
| FtsQ-Avi-R | ACTCGATCTTCTGGGCCTCGAAGATGTCGTTCAAGCCTCTAGATTGTTGTTCTGCCTGTGCCT | |
| LacZ-Avi-V-F | TTCCGGCTTGAACGACATCT | Cloning of the vector bone of plasmid pJH29-LacZ-Avi from pJH29-EspP-Avi with primer 29-V-R |
| LacZ-Avi-F | ATGTGGAACTGGCGGACTAATATGACCATGATTACGGATTCAC | Cloning of the *lacZ-Avi* tag segment |
| LacZ-Avi-R | AAGATGTCGTTCAAGCCGGAACCTTTTTGACACCAGACCAACTG | |
| pJH30-F | ATCAGTAAGTTGGCAGCATC | Cloning of the vector bone of plasmid pJH30 |
| pJH30-R | ATGGAGAAACAGTAGAGAGTTG | |
| S10 leader-F | CAACTCTCTACTGTTTCTCCATGGCTACCTAACAATGCTCC | Cloning of the S10 leader containing SD from MG1655 genome (wild-type SD) or MY1901 genome (mutant SD) |
| S10 leader-R | AAAGTTCTTCTCCTTTACTCATCGGATACGGATTCTTTGGTTCTG | |
| GFP-F | ATGAGTAAAGGAGAAGAACTTTTC | Cloning of the *gfp* segment |
| GFP-R | TGATGCTGCCAACTTACTGATCTATTTGTATAGTTCATCCATG | |
| pJH31-R | CGCGGAACCCCTATTTGTT | Cloning of the vector bone of plasmid pJH31 with primer pJH30-F |
| $P_{S10}$-F | AATAAACAAATAGGGGTTCCGCGGAGCGTGTCAAAAATGCACTG | Cloning of the $P_{S10}$ promoter and S10 leader containing SD from MG1655 genome or MY1901 genome |

**Supplementary Table S2. Summary of single nucleotide variants (SNVs), and insertions and deletions (INDELs).** The strain MY1901 shared common mutations with the original strain MG1655 in the laboratory, except the mutation located in the Shine-Dalgarno (SD) sequence of ribosome S10 operon. The evolved strain MY1901 by sub-culturing showed new mutations.

| Strain | Mutation site | Type | Chrom Start | chrom END | Ref | Obs | Relative to original strain |
|---|---|---|---|---|---|---|---|
| MY1901 | *fhuA* | SNV | 168123 | 168123 | A | C | |
| | *ybhJ* | SNV | 803662 | 803662 | C | A | |
| | *mntP* | SNV | 1905761 | 1905761 | G | A | |
| | *yjbI* | SNV | 4251856 | 4251856 | T | C | |
| | *gatC* | INDEL | 2173363 | 2173364 | CC | - | |
| | *glpR* | INDEL | 3560455 | 3560455 | - | G | |
| | *Promoter yagEp3* | SNV | 282234 | 282234 | G | T | |
| | *A repetitive extragenic palindrome (REP) element downstream of yjcO* | INDEL | 4296381 | 4296381 | - | GC | |
| | *Shine-Dalgarno (SD) sequence of ribosome S10 operon* | SNV | 3453283 | 3453283 | C | T | unique |
| MY1901 (evolved) | *rpsJ* | SNV | 3453146 | 3453146 | A | C | |
| | *DnaA* | SNV | 3882819 | 3882819 | G | T | |
| | *rpoB* | SNV | 4182952 | 4182952 | G | A | |
| | *rpoC* | SNV | 4186367 | 4186367 | C | A | |

Ref = reference allele; Obs = observed allele.

**Supplementary Table S3. Growth rate and the time cost of initiation steps ($T_{init}$) of cells under different nutrient conditions.** Growth rates were calculated from the corresponding growth curves that were shown in Supplementary Figure S4. All values are expressed as the mean ± standard deviation of the mean of samples. The initiation time ($T_{init}$) was calculated as described in the Materials and Methods section.

| **Strain** | **Nutrient condition** | **Growth rate ($h^{-1}$)** | **$T_{int}$** |
|---|---|---|---|
| MG1655Δ*lacZ* | Glucose + cAA<br>0.2% glucose + 0.2% casamino acids | 1.10 ± 0.04 | 10.0±0.2 (FusA) |
| | | | 9.6±0.2 (MsbA) |
| MG1655Δ*lacZ* | Glycerol + NH4Cl<br>0.2% glycerol + 10 mM $NH_4Cl$ | 0.55± 0.03 | 9.6±0.2 (FusA) |
| | | | 9.1±0.2(MsbA) |
| MY1905Δ*lacZ*Δ*cat* | Glucose + cAA<br>0.2% glucose + 0.2% casamino acids | 0.55± 0.02 | 11.7±0.3 (FusA) |
| | | | 10.2±0.2(MsbA) |

**Supplementary Table S4. The translation rates of FusA-LacZ and MsbA-LacZ under different growth rates.** The translation rate equals translation rate (aa $s^{-1}$) /336 (proteins $mRNA^{-1}$ $s^{-1}$) (Zarai et al., 2014), as the average length of proteins in *E. coli* is 336 amino acids (Gong et al., 2008). The elongation rate equals elongation rate (aa $s^{-1}$) /11 (sites $s^{-1}$) (Zarai et al., 2014) , as each ribosome occupies about 11 residues in *E. coli* (10). All values are expressed as the mean ± standard deviation of the mean of samples.

| **Strain** | **Growth rate ($h^{-1}$)** | **Translation rate (proteins $mRNA^{-1}$ $s^{-1}$)** | **Elongation rate (sites $s^{-1}$)** | **Initiation rate (sites $s^{-1}$)** |
|---|---|---|---|---|
| MG1655Δ*lacZ* | 1.10 ± 0.04 | (FusA) 0.047 ±0.0003 | 1.47±0.01 | 0.049±0.0003 |
| | | (MsbA) 0.045 ±0.0003 | 1.40±0.01 | 0.047±0.0003 |
| MG1655Δ*lacZ* | 0.55± 0.03 | (FusA) 0.040±0.001 | 1.24±0.04 | 0.042±0.001 |
| | | (MsbA) 0.038±0.001 | 1.15±0.03 | 0.039±0.001 |
| MY1901Δ*lacZ*Δ*cat* | 0.55± 0.02 | (FusA) 0.038±0.001 | 1.22±0.05 | 0.039±0.001 |
| | | (MsbA) 0.032±0.001 | 1.00±0.03 | 0.033±0.001 |

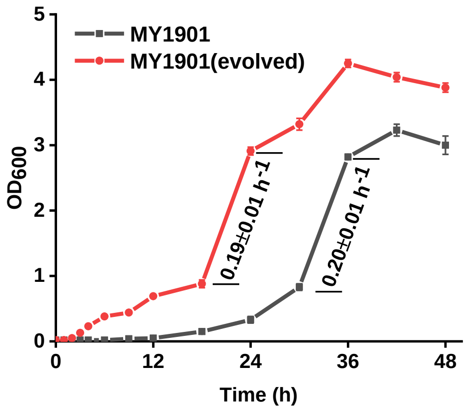


**Supplementary Figure S1** Growth curves of strains MY1901 and its evolved strain. Solid curves are the mean of three independent measurements and error bars represent the standard deviation of the mean of samples. All growth rates shown represent the mean ± standard deviation of three independent experiments.

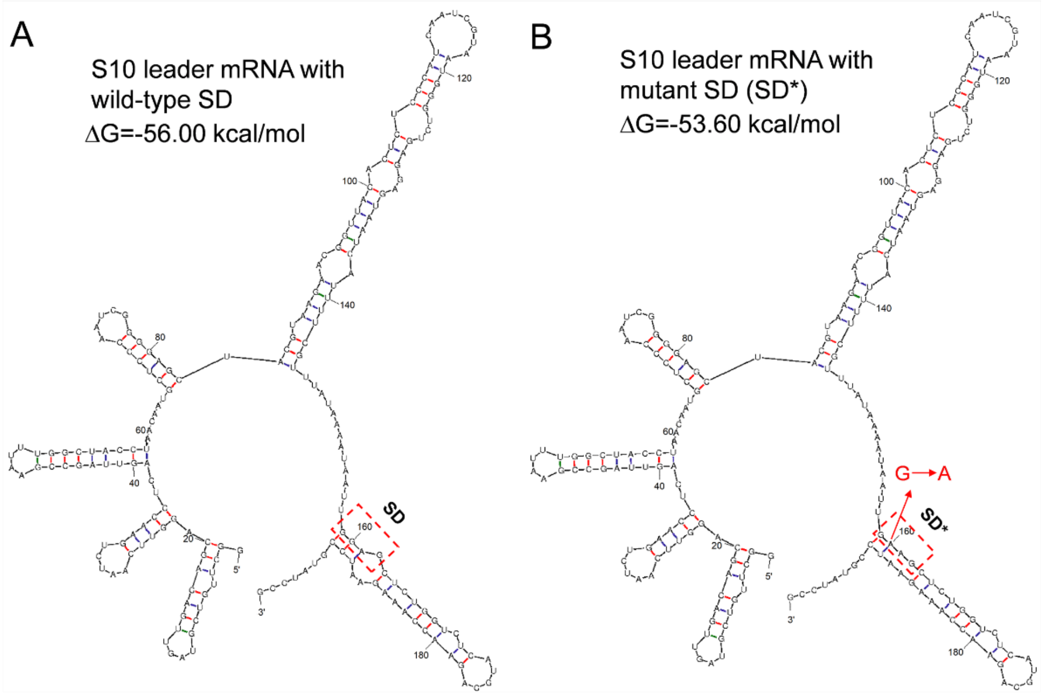


**Supplementary Figure S2** Predicted mRNA structures of the S10 leader. **(A)** RNA folding predicted hairpin structure of S10 leader with wild-type SD and the minimum free energy (MFE) of the thermodynamic ensemble (ΔG). **(B)** RNA folding predicted hairpin structure of S10 leader with mutant SD (SD*) and the MFE and the ΔG. The boxed region corresponds to the SD sequence.

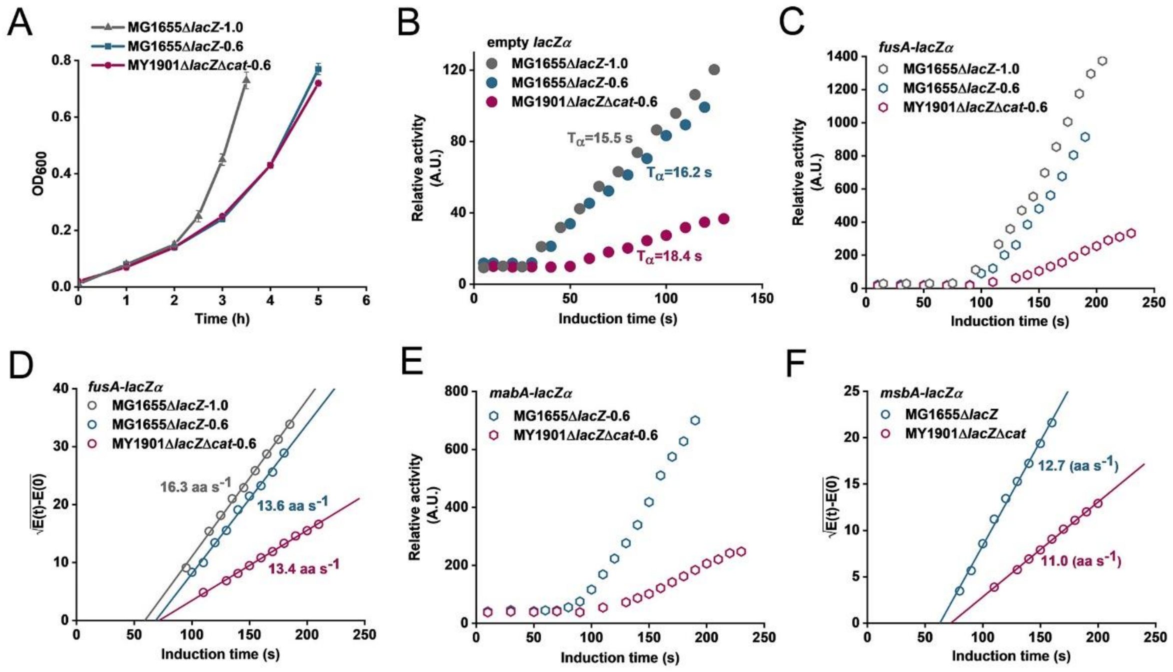


**Supplementary Figure S3** Measurement of translational elongation rates of wild-type and suppressor cells. **(A)** Growth curves of strains in different MOPS medium. The growth rates of MG1655Δ*lacZ* and MY1901Δ*lacZ*Δ*cat* grown in the Glucose + cAA medium were approximately 1.0 $h^{-1}$ and 0.6 $h^{-1}$, respectively. The growth rate of MG1655Δ*lacZ* grown in Glycerol + $NH_4Cl$ medium was approximately 0.6 $h^{-1}$(Supplementary Table S2). Solid curves are the mean of three independent biological replicates, and error bars represent the standard deviation value. **(B)** Calibration of the time cost of initiation steps by measuring the induction kinetics of the empty LacZα fragment. **(C)** The induction curves of the LacZα fused protein FusA-LacZα. **(D)** The Schleif plot of the FusA-LacZα protein was plotted against the induction time. **(E)** The induction curves of the LacZα fused protein MsbA-LacZα. **(F)** The Schleif plot of the MsbA-LacZα protein was plotted against the induction time. Schleif plots were repeated three times with one typical result shown here.

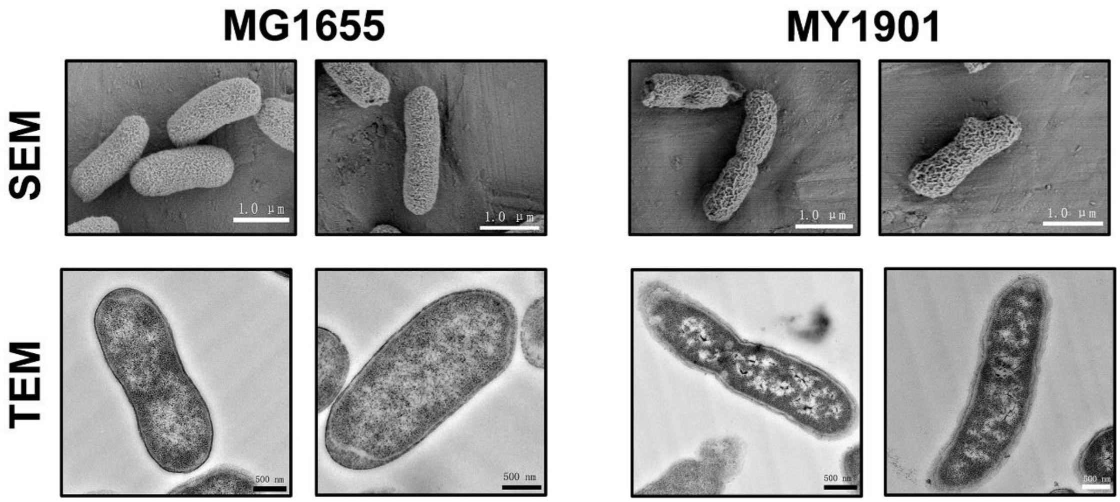


**Supplementary Figure S4** Scanning electron microscopy (SEM) and transmission electron microscopy (TEM) analysis of the wild-type strain MG1655 and the suppressor strain MY1901. For SEM, the scale bar is 1.0 µm. For TEM, the scale bar is 500 nM.

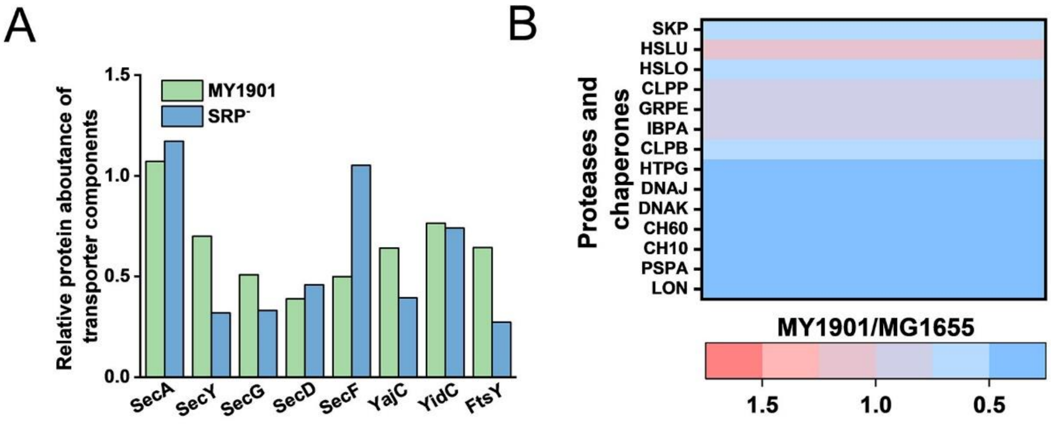


**Supplementary Figure S5** Fold changes in the expression of the components of transport machinery **(A)** and the heat shock response-related proteins **(B)** in strain MY1901relative to that in strain MG1655 (**Supplementary Data Set S1B** and **E**).
